# Uniform annotation framework reveals genome size and LINE/LTR retrotransposons as predictors of gene family expansion across Coleoptera

**DOI:** 10.64898/2026.03.25.714136

**Authors:** Milena Trabert, Jesper Boman, Elina Immonen

## Abstract

Gene family evolution – the turnover of duplicated homologous genes – shapes genome architecture and fuels phenotypic innovation. Repetitive elements (REs) facilitate gene duplication and genome expansion, yet whether variation in repeat abundance and genome size (GS) scales with gene family evolution across species remains unclear. Coleoptera provides a well-suited system for examining these dynamics because of its major ecological diversification and extensive genome size variation. Comparative tests of these relationships are however hindered by heterogenous genome annotations that distort gene counts and orthogroup assignments. We first evaluate how repeat- and gene-annotation strategies influence gene- and orthogroup-detection across beetle genomes. We then apply unified re-annotations of both to identify rapidly evolving gene families and test whether GS and repeat content covary with gene family size evolution. Nearly 500 orthogroups are rapidly evolving in Coleoptera, many of which are linked to ecologically crucial functions such as chemosensory perception and detoxification. GS and RE abundance are correlated, and on average scale positively with gene family sizes. LINE and DNA transposable elements commonly flank rapidly expanding gene families, but with pronounced species-specific variation. Together, these findings position genome architecture and repeat dynamics as fundamental determinants of gene family evolvability.

## Introduction

Gene families are groups of ancestrally related genes that evolve through gene duplication – a major mechanism that creates new genetic material and opportunities for evolution (Ohno 1970)(Lynch and Conery 2000). Classic examples of gene family expansions include the *Hox* genes, which pattern body plans across metazoans (Lemons and McGinnis 2006)(Holland 2013), and the Major Histocompatibility Complex (MHC) genes central to vertebrate immune defence (Fortier and Pritchard 2025). Following duplication, most new copies are eventually lost, restoring the ancestral single-copy state (Lynch and Conery 2000). However, several evolutionary trajectories can promote the retention of duplicates: neofunctionalization, where one copy acquires a new function; subfunctionalization, where ancestral functions are partitioned between copies; hypofunctionalization, involving reduced expression from both copies, maintaining the ancestral level; or dosage conservation, when both copies maintain ancestral expression, increasing the overall expression level (Birchler and Yang 2022). Duplications occur repeatedly along a phylogeny, interspersed with speciation, giving rise to homologous multi-member gene families shared across species (i.e. orthogroups, Fig 1A). Gene duplications that alter dosage or function to generate phenotypic variation within (Kaufmann et al. 2023)(Singleton et al. 2003) and between species (Davidson and Moczek 2024)(Dort et al. 2024) contribute to ecological diversification (Karunarathne et al. 2025), and ultimately speciation (Lynch and Force 2000).

**Figure 1:**
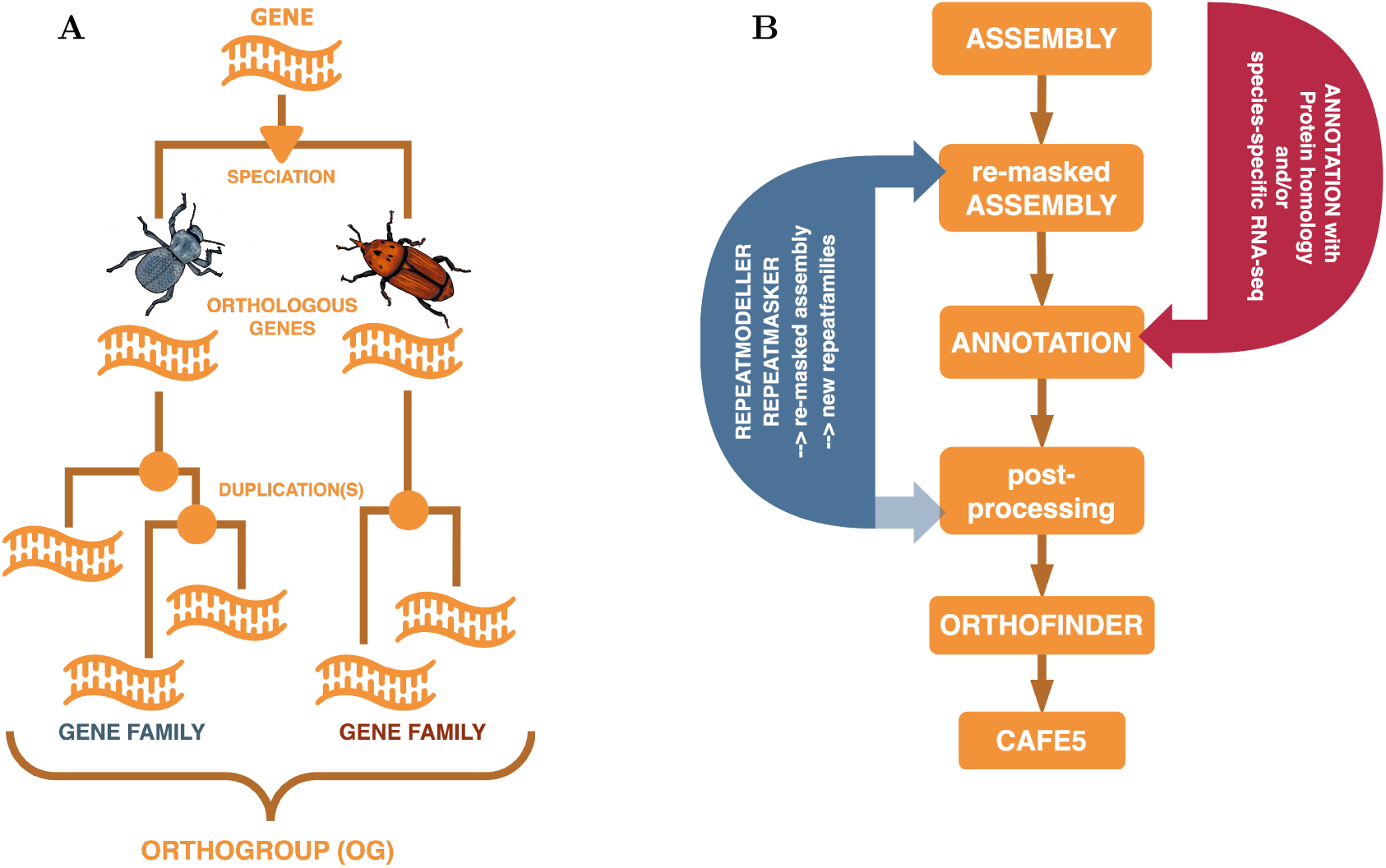
Schematic visualizations of key concepts. (**A**): Visual representation of the difference between gene family and orthogroup. This is one possible scenario, the speciation and duplication events can occur in any order. (**B**): Flowchart of the computational pipeline used to create the uniform annotations. Highlighted are the aspects we have varied in different ways to explore how they affect genome annotation. (blue) All assemblies we have used were already repeatmasked, but we have masked them again with a de novo generated repeat library (referred to as re-masking) and used the repeat libraries for an additional filtering step. (red) Genome annotation can be supported by protein evidence (from a wide variety of species) and also RNA-seq evidence (species-specific, higher quality evidence than protein), and we test how these data impact genome annotation in different ways.

Gene duplications can be mediated by repetitive elements on a genome-wide or localized level. On a genome-wide level, repeats contribute to structural variation by promoting double-strand breaks and nonallelic homologous recombination, leading to duplications, deletions, and rearrangements (Bose et al. 2014)(Reams et al. 2012) (reviewed in (Reams and Roth 2015)). In insects, repeat abundance varies strikingly among genomes, from ∼10% in the Antarctic midge *Belgica antarctica* (99Mb genome; Diptera (Kelley et al. 2014)) to ∼70% in the seed beetle *Callosobruchus maculatus* (Arnqvist et al. 2024) (1.2GB genome; Coleoptera (Arnqvist et al. 2015)). Genomes also differ in TE composition, including the relative abundance of LINEs, LTRs, DNA transposons (e.g. Helitrons/pack-MULE-like) and SINEs (Petersen et al. 2019a). This composition is functionally important: LINEs generate intronless retrogenes, Helitrons and pack-MULE–like elements mobilize gene fragments or entire genes, and abundant LTRs and other repeats act as substrates for non-allelic homologus recombination that duplicate nearby loci (Tan et al. 2021) (reviewed in (Cerbin and Jiang 2018) and (Lallemand et al. 2020)). Consequently, TE-rich genomes provide multiple structural pathways for generating new paralogs, potentially accelerating gene-family expansions and turnover (Cerbin and Jiang 2018). However, the long-term retention of duplicates is shaped by selective and functional constraints — such as dosage balance, network connectivity, and purifying selection — which can decouple gene family dynamics from repeat proliferation. Thus, while TEs provide a potent source of structural novelty, their genome-wide contribution to gene family evolution remains an open question.

In many taxa, there is a positive association between repetitive element content of the genome and the total genome size (GS) (Petersen et al. 2019b). Larger genomes may provide more opportunities for gene duplication and the subsequent evolution of gene families, directly or indirectly in association with repetitive elements. Genome size is a fundamental biological characteristic, and its evolutionary causes and consequences are topics of ongoing debate (Petrov et al. 2000)(Kapusta et al. 2017)(Gregory 2004)(Blommaert 2020). Eukaryotic organisms vary enormously in GS, ranging from 2.9Mb in the intracellular parasite *Encephalitozoon cuniculi* (Vivarès and Méténier 2004), over 3.1Gb in humans (Piovesan et al. 2019), to 160Gb in the Caledonian fork fern (Fernández et al. 2024), but there is no clear relationship between GS and organismal complexity among eukaryotes. Closely related species (Albach and Greilhuber 2004)(Hospodářská et al. 2025) and even populations (Boman and Arnqvist 2023) show considerable variation in genome size. However, while GS and gene number (GN) scale positively across Eukaryota (Elliott and Gregory 2015), GN varies much less than GS. Comparative analyses that jointly examine genome size, repeat content, and gene family evolution are lacking, leaving the interplay among these genomic features still poorly resolved.

While the increasing availability of genome assemblies has enabled large-scale analyses of gene family dynamics, accurately determining gene content remains a non-trivial and often underappreciated technical challenge. The same TE burden that biologically contributes to gene birth, also complicates genome assembly, repeat masking, and gene prediction, and may lead to missed, fragmented, or TE-derived gene models that distort inferred family sizes (Saenko et al. 2025) (Weisman et al. 2022)(Dort et al. 2024). Annotation methods differ in how genes are identified (Weisman et al. 2022), which can bias orthologue inference and inflate lineage-specific gene counts (Van Oss and Carvunis 2019)(Dort et al. 2024). These problems are exacerbated in repeat-rich genomes, where assembly contiguity strongly influences repeat detection: highly contiguous, long-read assemblies tend to recover more repeats (Sproul et al. 2023)(Peona et al. 2021), and differences in repeat reference databases further affect gene prediction outcomes through repeatmasking (Bayer et al. 2018). Inadequate or inconsistent repeatmasking can therefore generate both false-positive gene models derived from TEs and false negatives where genuine genes are obscured within repetitive regions (Elsik et al. 2014), leading to biased cross-species comparisons. Although these issues can be partially mitigated by re-annotating genomes using a uniform gene prediction pipeline (Saenko et al. 2025), uniformity in gene prediction alone may be insufficient without a consistent repeat annotation strategy. Moreover, the common practice of genome annotation liftover between assemblies of differing contiguity or repeat resolution, with tools such as liftoff (Shumate and Salzberg 2021), may further propagate these biases. Together, these effects underscore that heterogeneity in assembly quality, repeat annotation, and gene prediction pipelines can substantially confound comparative genomic analyses, yet remain insufficiently discussed. Recent demonstrations of annotation heterogeneity impacts on lineage-specific gene, and ortholog inference (Weisman et al. 2022) (Prieto-Baños et al. 2025), exemplify this challenge.

Here, we investigate gene family evolution and putative connections with genome size and TE abundance in the order *Coleoptera* (beetles). Despite being regarded as the most species-rich animal order, *Coleoptera* are underrepresented in genome sequencing projects (Hotaling et al. 2021), and little is known about gene family and repetitive element evolution in this order. Beetles show extensive ecological and reproductive diversification, and pronounced variation in genome size (e.g. 204 Mb in *Tribolium castaneum* (Brown et al. 1990) to 1.2 Gb in *Callosobruchus maculatus* (Arnqvist et al. 2015)), making them a very interesting group to assess genome-wide relationships between gene family expansions, repetitive elements (REs) and GS. We address these in a comparative analysis of 13 species across Polyphaga, the largest suborder of Coleoptera, spanning over 250 Mya and containing more than 300,000 described species (Zhang et al. 2018)(Slipinski et al. 2011).

First, we address how genome annotation strategies affect gene prediction, completeness metrics and protein length distribution. In combination with phylogeny-aware methods we show the effect of annotation strategy on orthogroup and gene family sizes. We also demonstrate the impact of repeat annotation strategy, by comparing heterogenous and uniform repeat annotation approaches. Second, we characterize repetitive element landscapes across the species and test for correlation between RE abundance and GS. Third, by using orthogroups from uniform genome re-annotations and accounting for phylogenetic relationships, we identify rapidly evolving gene families in Coleoptera and investigate gene family size variation in relation to GS and RE abundance. Our comparative framework reveals that GS and RE abundance are positively correlated and consistent predictors of average gene family size across beetle genomes. Nearly 500 gene families show signatures of rapid expansion, reflecting divergence in ecologically crucial functions such as chemosensory perception and detoxification.

## Results

### Comparison of genome annotation strategies

Our comparative dataset includes 13 Coleoptera genomes across the suborder Polyphaga that span four superfamilies (Elatoeroidea, Coccinelloidea, Tenebrionoidea, Chrysomeloidea), six families and twelve genera (species included: *Acanthoscelides obtectus*, *Asbolus verrucosus*, *Bruchidius siliquastri*, *Callosobruchus chinensis*, *Callosobruchus maculatus*, *Coccinella septempunctata*, *Dendroctonus ponderosae*, *Ignelater luminosus*, *Photinus pyralis*, *Rhynchophorus ferrugineus*, *Tenebrio molitor*, *Tribolium castaneum*, *Zophobas morio*), with *Drosophila melanogaster* included as the outgroup. We generated uniform genome annotations for each of the species (using the *ab initio* and homology-based hybrid strategy implemented in BRAKER3). We compared these to the available published annotations on the same genome assemblies (hereafter “native annotations”) to assess the consequences of gene annotation heterogeneity. The BUSCO scores in the two types of annotations are high and largely similar (Fig. 2A), demonstrating robust prediction of conserved BUSCO genes. The annotations do not differ in predicting missing or duplicated proteins either, suggesting that these flaws are due to assembly quality and not affected by annotation choices investigated here. However, some genomes show drastic differences in the predicted gene count between the annotations (Fig. 2B).

**Figure 2:**
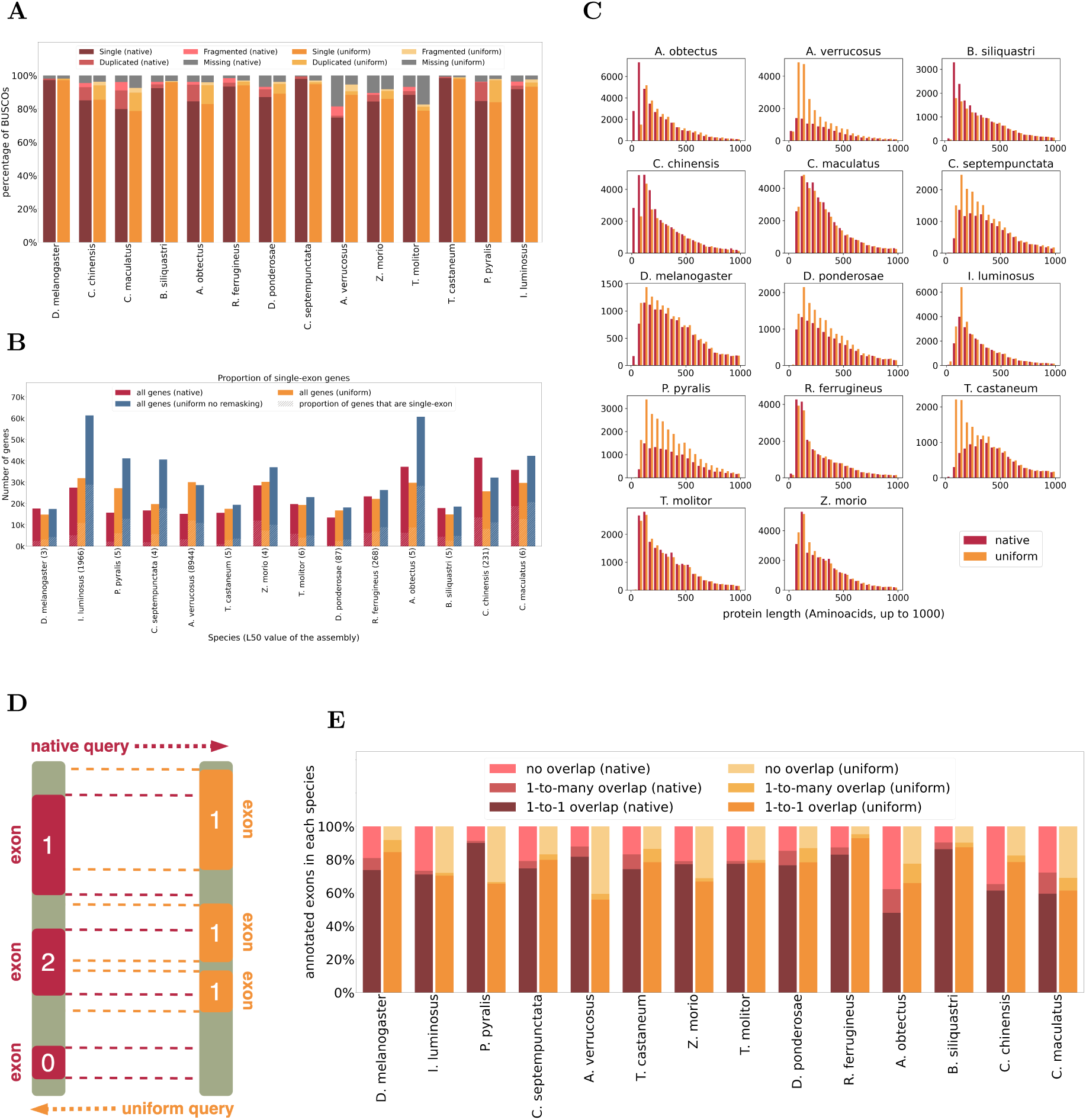
Evaluation of the uniform annotation pipeline. (**A**): BUSCO evaluation of both annotation sets, red bars are the native annotation, and orange are the uniform annotation. For most assemblies the scores are very similar. (**B**): Number of genes annotated in three different annotation methods and proportion of which are single-exon. Red bars are the heterogeneous native annotations, yellow bars are the homogeneous annotations as described in Fig. 1B, and the blue bars are also the uniform annotation pipeline, but without uniform repeatmasking. (**C**): Histograms of protein length distribution for all species in both annotations. Native annotations (red) show variable shapes of the histograms, while uniform (orange) annotations show similar distributions in all assemblies. Note that all species are only shown up to 1000 AAs. (**D**): The two annotations are based on the same assembly, therefore their annotated features can be compared for position overlap. Visual representation of calculating the overlap of two annotations based on the same assembly. Each annotation is used as a query once, and for every exon in the query annotation the number of exons it overlaps with in the reference annotation are counted. (**E**): For each species, the overlap of annotated exons with native query (red) and uniform query (orange) is shown. A feature can overlap with features in the other annotation in three ways: 1-to-1, 1-to-many (multi-overlap), or no overlap. The number of exons that show 1-to-1 overlaps is the same for both annotations in a species, but this number makes up a different proportion of the total number of annotated exons for each annotation method.

To understand such discrepancies further, we examined repeat annotation as a source of annotation heterogeneity. Comparing uniform annotations with and without assembly re-masking (Fig. 2B) shows that in several cases re-masked assemblies yield substantially lower gene number (GN) compared to the non-remasked, which are also closer to the native annotations. The large differences in GN for many species strongly suggests that repeats have been insufficiently masked and become falsely identified as “host” genes. False-positive TE genes in gene annotations have the potential to cause serious misconceptions about the evolution of both genes and repeats. The inadequate masking here is greater in some assemblies than others, but does not systematically increase with genome size (spanning ∼400 Mb to ∼1 Gb in the three worst cases) or assembly fragmentation (see assembly L50 values in Fig. 2B). The proteins lost after re-masking are mainly very short (See Fig. S4), and non-re-masked assemblies show an elevated proportion of single-exon genes, indicating that especially short proteins are false-positives caused by inadequate repeat masking.

Hereafter we only consider the uniform annotations based on the complete re-annotation pipeline (including re-masking and repeat consensus sequence filtering, see Methods). The protein length distributions in the native annotations differ substantially between the species, whereas the unified annotations show highly consistent patterns of variation (Fig. 2C). The native and uniform annotations differ mainly in the abundance of proteins shorter than 500 amino acids, with no consistent directionality. As the uniform annotations are based on the same assembly as the native annotations, we can compare the location of individual exons in both annotations for each species (Fig. 2D-E). The 1-to-1 position overlap for a pairwise comparison is achieved in about 80% of the cases across most species for individual exons (less for whole transcripts, see SR5). The vast majority of 1-to-many overlaps in either direction are 1-to-2 overlaps (Fig. S6-7), with the transcript-based comparison showing slightly more 1-to-3 overlaps (Fig. S8-9). A 1-to-many overlap on the exon level means that the annotation struggles with correct exon-intron boundaries, while on the transcript level it is more indicative of genes being split or the wrong exons being grouped.

We also evaluated how RNA-data source generates heterogeneity in gene prediction when using evidence-based annotation strategy using RNA-seq data from three geographically distinct populations (Togo, Nigeria, and India), applied to a single focal genome assemly (of *C. maculatus* Lomé population from Togo). The three resulting annotations differ in detected gene number by over 800, with the number of genes decreasing with geographic distance (Lomé RNA: 15,703 genes, Nigeria RNA: 15,496 genes, and South-India RNA: 14,881 genes, see Figures S1 and S2)). Furthermore, each RNA-seq source generated 7-13% unique genes not predicted in the other two annotations. Taken together, these analyses demonstrate how RNA-seq source and insufficient repeat masking introduce substantial variation in gene prediction. Re-masked uniform annotations match native annotations in conserved gene recovery while reducing annotation heterogeneity and false positives.

### Repeat abundance and genome size variation

Upon establishing uniform gene- and repeat-annotations we can now study genome evolution in Coleoptera at an unprecedented level of rigor and comparability. First, we investigated the relationship between repeats and genome size. The repeat libraries generated for all species reveal considerable variation in the repeat content (Figure S10). The total abundance of repetitive elements ranges from ∼20% in *D. ponderosae*, to ∼75% in *C. maculatus* and *A. obtectus*. Generally, Retroelements and DNA transposons are the most common repeat categories together with the unclassified repeats, but which class dominates varies even between closely related species. As expected, the proportion of unknown repeats is near 0% in the Diptera outgroup (*D. melanogaster*) but considerably higher in all beetle species, due to the fact that *D. melanogaster* is well represented in repeat reference databases compared to non-model insects (Sproul et al. 2023). Within each genome, the repeat abundance also varies depending on chromosome location (see Supplementary Figures Section 1 for repeat abundance across the genome for each species).

We tested for an association between genome size (GS) and genome-wide repeat content using GS estimates obtained from genome assemblies that correspond closely to flow cytometry estimates (Pflug et al. 2020). GS varies by six-fold across the species, ranging from ∼200 Mb to 1.2 Gb (see Fig. 3A). As predicted, the genome size and total repetitive element abundance are significantly positively correlated across the phylogeny (linear regression using phylogenetically independent contrasts (PIC), p=0.0026, F-statistic = 10.13).

**Figure 3:**
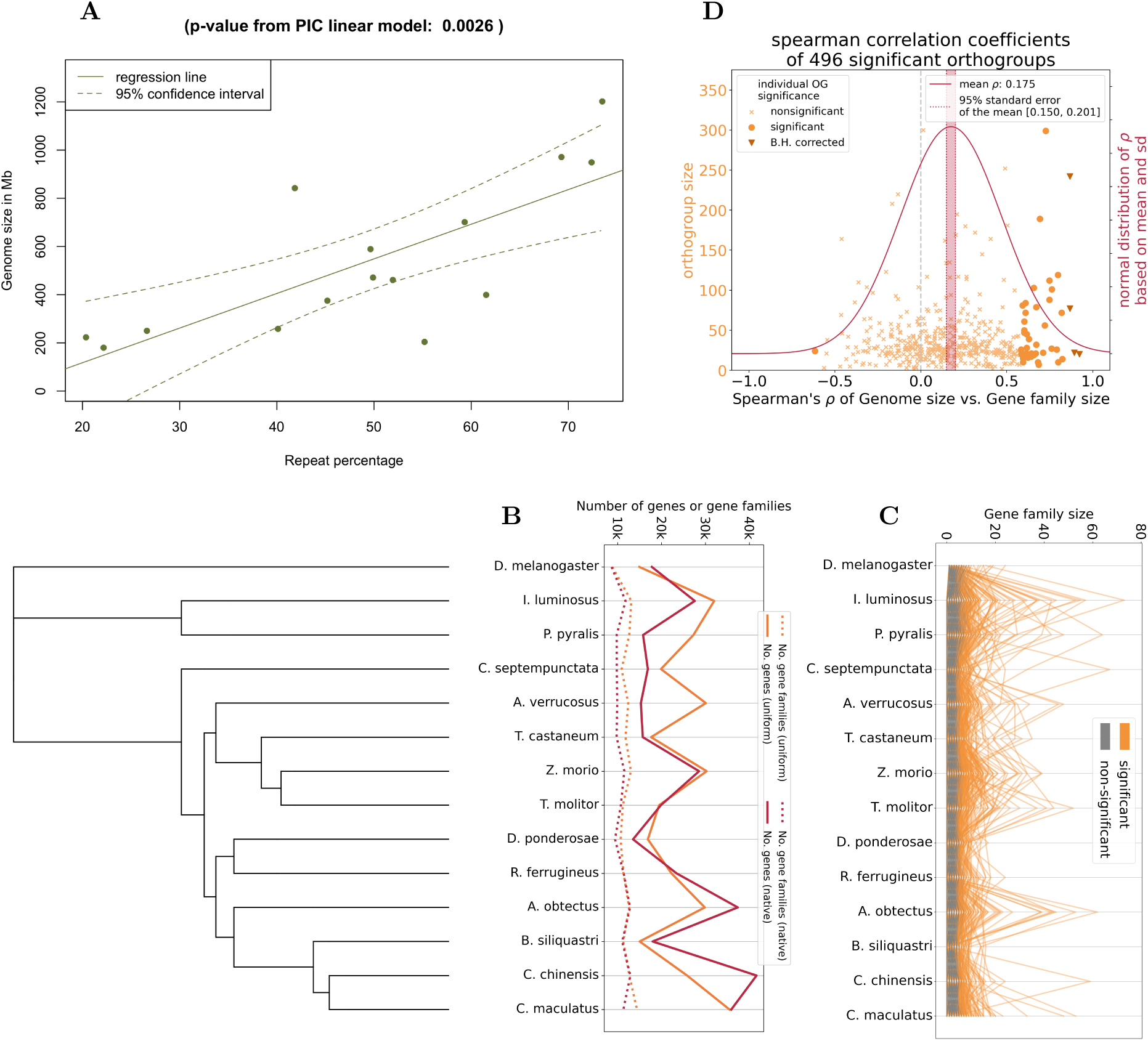
Orthofinder and CAFE5 results. (**A**): Scatterplot of repeat content vs. genome size of all species. The regression line and confidence interval are based on a linear regression of the plotted values, but the p-value was calculated based on a linear regression with the phylogenetically independent contrasts (PICs) of both variables. (**B**): Number of orthogroups and all genes in orthogorups in each species as identified by Orthofinder (genes not assigned to an orthogroup are not included). (**C**): Number of members each orthogroup has in each species (gene family size). Separated into significantly rapidly evolving (orange) and not rapidly evolving (gray) according to CAFE results. Two very high peaks in *A. obtectus* have been identified as transposases and are therefore not shown and removed from further analysis (see SI section 5.2). (**D**): We have calculated the Spearman correlation coefficient for every orthogroup: within the orthogorup, every species is a data point with GS correlated with species gene family sizes (for repeat content instead of GS see Fig. SR12). All gene family sizes, repeat proportions, and genome sizes are transformed to phylogenetically independent contrasts. Orthogroups with individual significant p-values from the correlation are highlighted before and after Benjamini-Hochberg correction for multiple testing. Individual orthogroups cover a wide range of correlation coefficients, both positive and negative, but after multiple-testing correction none are significant with repeat content, and only very few for GS (Flybase orthologs: FBgn0017414, FBgn0028931, FBgn0032456 and FBgn0259199). The mean correlation coefficient is above 0.

### Gene family evolution

#### Gene family expansions are associated with repeats on a local genomic scale

We used Orthofinder to identify orthogroups, defined as all members of a homologous gene family across species (Fig. 1A). The number of orthologous gene families detected in each genome is highly consistent, at approximately 10k, while the total number of genes in all families varies substantially across species, in both annotation versions (Fig. 3B). We restricted the subsequent analyses to only highly conserved orthogroups with a member in *D. melanogaster* outgroup (present at the root of the tree) to test for significant gene family expansions (with CAFE5) across the phylogeny. Out of the 8,315 orthogroups analyzed, 496 (6%) are significantly rapidly evolving (see supplementary file 1). The significant orthogroups show large variation in gene family size between beetle families but in some cases even within genus (Fig. 3C). Species with many genes also tended to have larger gene family expansions.

We investigated putative associations between the 496 significant gene family expansions, genome-wide repeat content and GS, by estimating Spearman correlations. On average, gene family sizes and GS are positively correlated (Fig. 3D), with the mean correlation coefficient across all orthogroups significantly different from zero (based on non-overlapping standard error of the mean (SEM) around mean Spearman’s ρ = 0.175, SEM = [0.150, 0.201]). When considering individual orthogroups, correlations are significantly positive after multiple testing correction in four cases (Fig. 3D) (orthogroups with the following annotated *D. melanogaster* members: N0.HOG0000049, flybase ID FBgn0017414, *cag*, ρ = 0.867; N0.HOG0000296, flybase ID FBgn0028931, ρ = 0.867; N0.HOG0000761, flybase ID FBgn0032456, *Multidrug-Resistance like Protein 1, MRP*, ρ = 0.893; N0.HOG0002345, flybase ID FBgn0259199, *Snurprotin*, ρ = 0.923). Three of these orthogroups correlated with GS are directly or indirectly related to gene expression regulation. *Cag* is associated with DNA binding, FBgn0028931 is involved with regulation of RNA polymerase II, *MRP* is a transmembrane transport protein linked to toxin response and expressed during the entire lifespan, and *snurprotin* is involved in importing small RNAs into the nucleus. When testing for a correlation with gene family size and repetitive element abundance, the correlations are not individually significant for any orthogroup after multiple testing correction (Fig. S12). However, on average the correlation coefficients across all orthogroups are positive (mean ρ = 0.095, 95 % CI = [0.070, 0.119]), although weaker than for GS. In addition to using estimates of total repeat content, we estimated correlations for each separate repeat category (Fig. S13 and S14). After FDR correction, one expanding orthogroup shows a significant positive correlation with abundance of low complexity regions (Fig. S13B, orthogroup N0.HOG0000056 with ρ = 0.895). This orthogroup includes *D. melanogaster* ortholog *Odorant-binding protein 56e* (*Obp56e*), with peak expression in adult males. There is also a significant correlation between one orthogroup and satellite DNA content (Fig. S13D, *D. melanogaster* ortholog flybase ID FBgn0037955 *Kynurenine aminotransferase*, ρ = 0.965). Taken together, gene families tend to be larger in larger genomes, with a weaker but positive association with repeat content.

Gene duplication mechanisms differ in the genomic proximity of resulting duplicates: nonallelic homologous recombination generates tandem arrays, whereas retrotransposition inserts copies at dispersed genomic locations. While there is enrichment of intronless genes among rapidly evolving orthogroups (Fig S15), gene family size does not predict duplicate proximity across the phylogeny (Fig. S16), indicating that no single mechanism dominates gene family expansions.

The repeat landscape across scaffolds in each species show large variation along chromosomes, with centromeres and telomeres often being enriched with different repeat classes (Fig. S22-31, e.g. *A. obtectus* (S22) for telomeres and *B. siliquastri* (S23) for centromeres as prominent examples). To further probe the mechanisms that may duplicate individual genes and facilitate their persistence, we investigated the characteristic repeat sequences flanking the rapidly expanding gene families. For this, we compared repeat abundances within 10kbp up- and downstream of non-significant (hereafter referred to as the background) and significantly expanding gene families within a given genome (hereafter referred to as the foreground, with min. of two members per family) (Fig. 4A and C, all species in Fig. S32-45). The enrichment patterns were compared with a Wilcoxon test, by sampling the proportion of each repeat category every 500 bp to increase sampling independence (Fig. 4). In all species, the repeat content reduces drastically in the most proximate 1kb region flanking genes, as expected. DNA transposons, LINE retrotransposons and long terminal repeats (LTRs) are generally enriched around the foreground genes, especially in the closer proximity to the genes (1 to 4kb up- and downstream). This is a common theme regardless of their relative genome-wide abundance in a given genome (Fig. S10 and S22-31). However, the most common repeat categories in a given genome also tend to show larger effect size in the Wilcoxon test (Fig. S17 and S18). To complement the enrichment analysis of different repeat classes, we also tested whether the shape of the repeat landscapes for the classes significant in the Wilcoxon test differ between the background and foreground genes in each genome, using polynomial regression analysis (Fig. S46-103). The polynomial models support increasing differences between the foreground and background closer to the genes (based on 83% CI non-overlap, corresponding to ≈ p = 0.05) (e.g. Fig. S53 for *B. siliquastri* LINEs, or *R. ferrugineus* LINEs in Fig. S83), and in many species significant divergence over the entire examined length of the landscape (e.g *C. septempunctata* LINEs in Fig. S65, or *T. castaneum* LINEs in Fig. S89 among others). However, in cases where the repeat percentage of the focal class is generally low, the confidence intervals often overlap (e.g. *B. siliquastri* Rolling-Circles, Fig. S55; *C. maculatus* LINEs in Fig. S60 and LTRs in Fig. S61).

**Figure 4:**
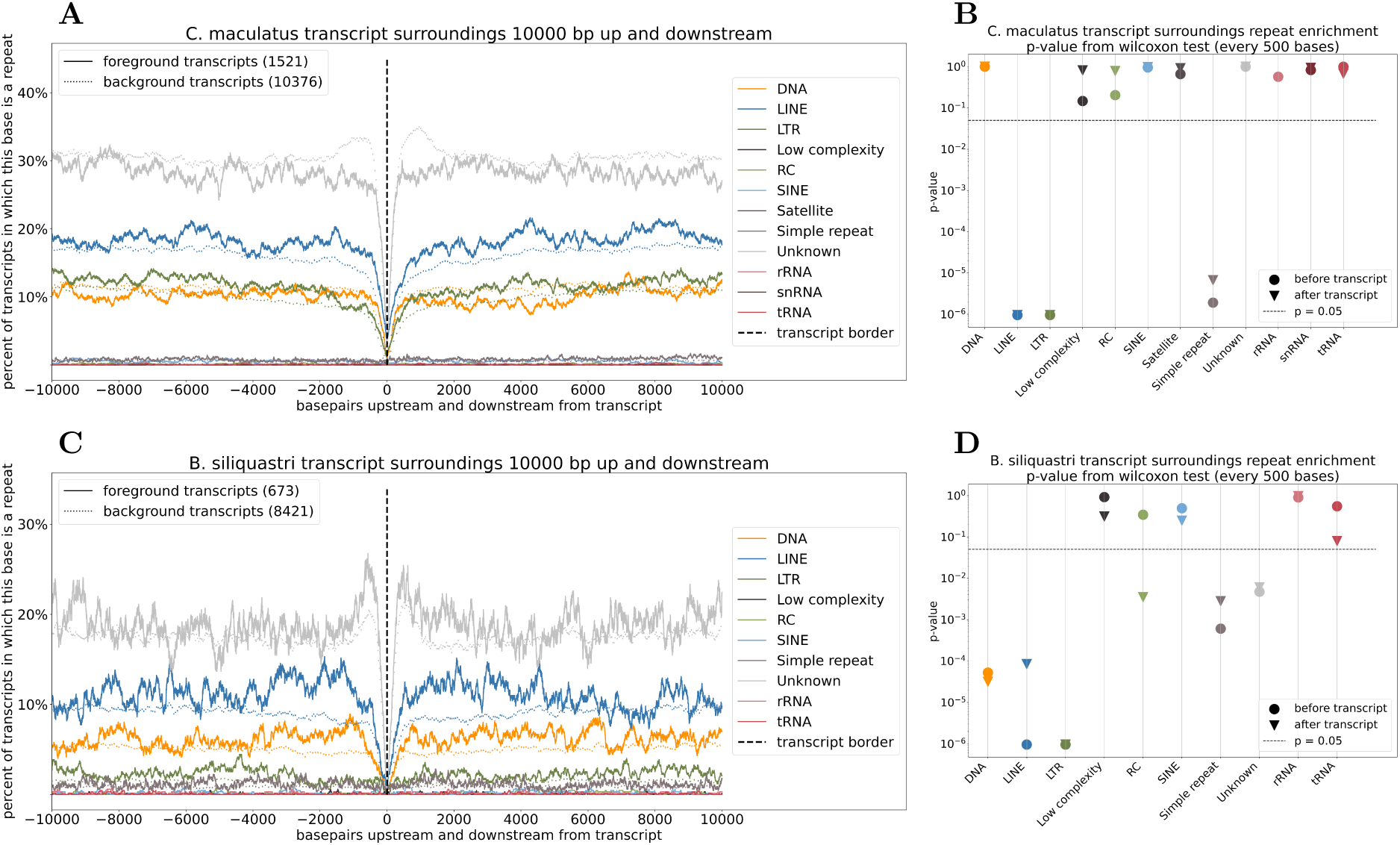
Repeat content in the surroundings of significantly expanding transcripts. (**A**, **C**): Abundance of different repeat classes according to repeatmasker annotations around transcripts that are part of significantly rapidly evolving orthogroups. All transcripts in an individual species are filtered to include only orthogroups that are actually expanding in this species (at least two gene family members) to only include transcripts potentially duplicated through TE activity. We show two exemplary species *C. maculatus* (**A**) and *B. siliquastri* (**C**), all species can be found in the Supplementary Figures section 2. All plots are based on the same number of significant and nonsignificant orthogroups from the CAFE5 analysis, but the gene family sizes within the orthogroups differ (see Fig. 1A), resulting in the different transcript numbers between species. (**B**, **D**): p-values form Wilcoxon tests for higher repeat content in expanding transcripts for *C. maculatus* (**B**) and *B. siliquastri* (**D**). Repeat content data was downsampled to only contain every 500th base, meaning 20 points before and 20 after the transcript, to increase the independence. See Supplementary Figures section 3 for Wilcoxon test results for all species, and plots of polynomial regressions of the significant repeat categories in all species.

We conclude that genome size is a stronger predictor of gene family expansions than the genome-wide repeat content, however, there are strong local associations. Significantly expanding gene families occur in genomic environments enriched especially with LINES, LTRs and DNA transposons, but the associations are also nuanced and specific to each species.

#### Coleopteran trait evolution is linked to gene-family expansion

When failing to sufficiently mask repeats, many rapidly expanding “gene families” are likely to be TEs and not genes of the host genome. Our uniform annotation, which included re-masking of repeats, has the potential to increase the true positive rates of significantly expanded gene families. With a higher true positive rate, a more relevant functional annotation can be made. To functionally characterize the rapidly expanding gene families in Coleoptera, we utilized the outgroup species *D. melanogaster* orthologs for functional annotation via FlyBase and clustering of significant orthogroups into functionally related groups using DAVID Bioinformatics tools (Sherman et al. 2022) and *a priori* information of categories known to be important or rapidly evolving in other insect orders and in the fireflies (*Elateriformia*) (Fallon et al. 2018a) (Supplementary File 1, Fig. 5A, Table S2). Twenty-three significantly rapidly evolving orthogroups, containing 1603 transcripts across all species, are functionally clustered with roles in chemosensory activity (Fig. 5B), including odorant binding proteins, olfactory receptors, ionotropic receptors, antennal transmembrane transporters and pheromone binding proteins (Table S3). Related to these, multiple orthogroups are also involved in pheromone synthesis (Figure 5C, Table S4). The gene family sizes of pheromone and general odorant binding proteins are largest and also show greatest and relatively concordant fluctuation in numbers across species, while the other chemosensory orthogroups show more independent gene family expansions in different species (Figure 5B). Other notable expansions have occurred in orthogroups related to detoxification in host adaptation (Figure 5D, Table S5), sexual reproduction and immunity (Figure 5E and Table S6). Further categories are presented in Figures S19-21 and corresponding Tables S7-9. We identify also an expansion in the gene family that contains luciferase, which causes fluorescence in *Elateriformia* (Fireflies) (Fallon et al. 2018a). Luciferase arose from Acyl-CoA synthetase, and this gene family and related ones show prominent expansions in the *Elateriformia* species *I. luminosus* and *P. pyralis* (Fig. S19 and Table SR7).

**Figure 5:**
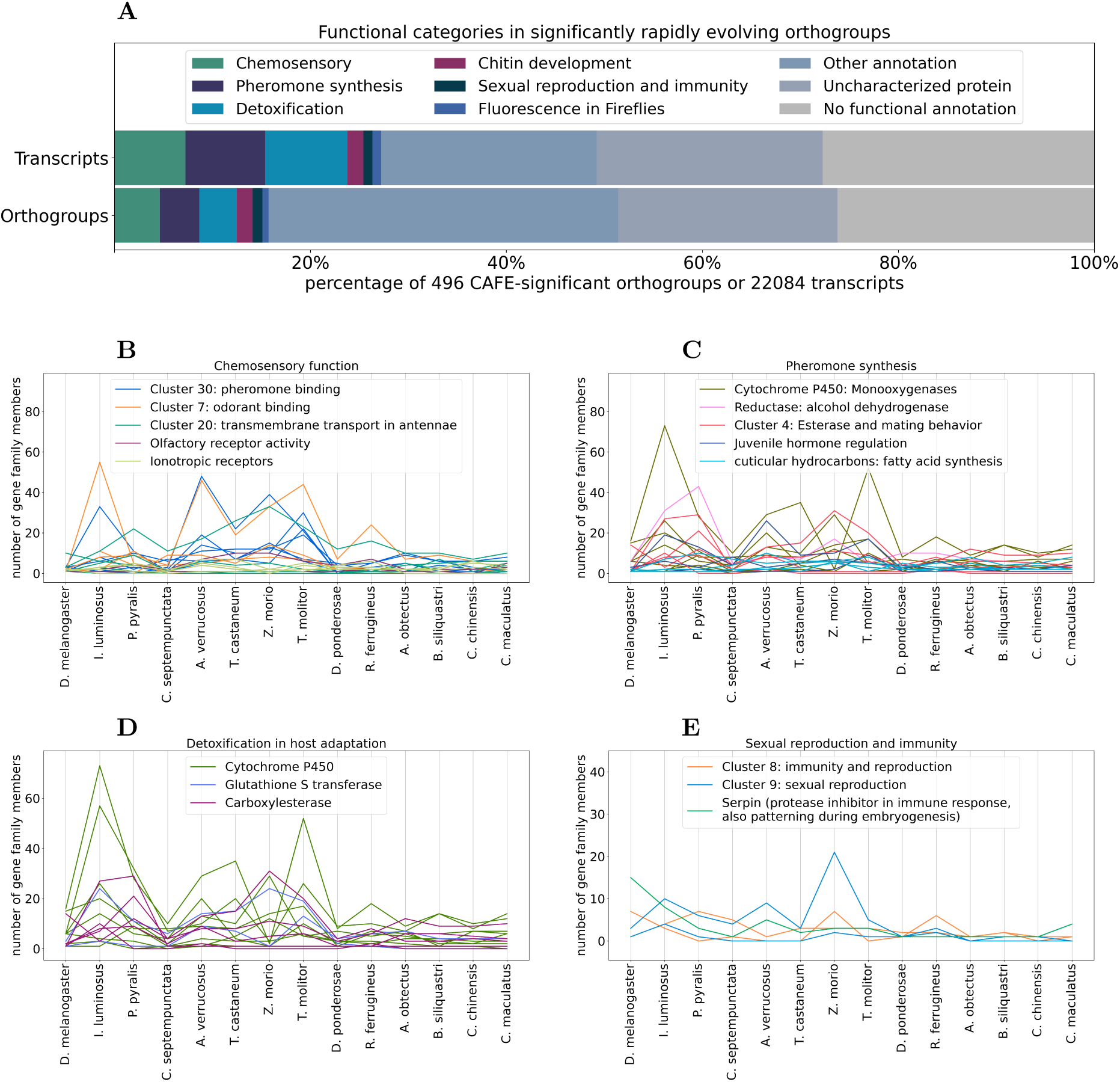
Functional evaluation of significantly rapidly evolving orthogroups. We have functionally clustered the significantly rapidly evolving orthogroups as described in the text and then evaluated them according to functional themes that have been found to be rapidly evolving in other insect orders. (**A**) We show the proportion of the identified functional themes in all the significantly rapidly evolving orthogroups. The proportion is higher when looking at the transcript numbers, compared to orthogroup numbers, showing that these functional categories are often in expanding gene families. (**B**, **C**, **D**, **E**) Gene family sizes of functional groups of interest. Each line is an individual orthogroup, showing the number of gene family members that orthogroup has in each species.

## Discussion

We develop and apply a uniform genome annotation pipeline for comparative genomics based on publicly available data and compare the resulting uniform annotations to preexisting heterogeneous native annotations. We find that the uniform annotations eliminate sources of heterogeneity while not compromising quality on the genome-wide scale. Further, we identify a large variability in gene number and gene family size across the studied species in addition to consistent patterns linking gene family evolution, genome size (GS) variation and repetitive element (RE) abundance. On average, both GS and RE content show positive associations with rapidly expanding gene family sizes. Nearly 500 ancestral gene families are significantly rapidly evolving, and they include functions related to chemosensory perception and detoxification – traits central to ecological adaptation in Coleoptera. Together, our results provide methodological insights into annotation biases commonly overlooked in comparative genomic analyses, and clarify how genome architecture can shape evolutionary processes acting on gene families.

### Annotation strategy and methodological implications

Our analyses first show that genome annotation strategy is a major, yet often underappreciated, source of bias in comparative genomics. While work that takes the bias caused by heterogeneous annotation methods into account has started to emerge (Weisman et al. 2022)(Prieto-Baños et al. 2025), the field is still far from having a consensus on the use of heterogeneous publicly available data. Differences in the provenance and composition of RNA-seq evidence—even when applied to the same assembly—alter gene predictions, orthogroup assignments, and protein length distributions. Such heterogeneity is common in publicly available annotations because both related transcriptomic data typically vary in population origin, developmental stage, or environmental context, and genome annotation pipelines are not standardized. These factors directly affect which transcripts are represented in gene models. As demonstrated previously by Weisman et al. (2022) and Dort et al. (2024), non-uniform annotation pipelines can artificially inflate lineage-specific gene counts and distort estimates of gene family turnover. Our results extend these findings by showing that even within a single assembly, heterogeneous RNA evidence produces inconsistent annotations. When using RNAseq-evidence-based annotation pipelines for comparative genomics analysis, the RNAseq data for all species should therefore be from the same population as the assembly, and of comparable quality both between species and between samples of the same species, which is very difficult to achieve when relying on publicly available data. In Figure 2, we compare the uniform (RNAseq-less) annotations to native (RNAseq-based) annotations and conclude that there are often major structural differences. Therefore, the heterogeneity introduced by using RNA-seq based annotations together with RNAseq-less annotations in the same comparative study would be a major source of technical variation, likely biasing the results. In addition, we show the importance of uniform repeatmasking. When the annotations are not uniformly repeatmasked, up to 20k surplus genes are predicted compared to when repeatmasking is re-applied consistently (see Fig. 2B). Thus, lack of uniform repeatmasking can lead to erroneous conclusions on the evolution of genome architecture. Repeatmasking depends on the pipeline used to predict repeat consensus sequences (also referred to as repeatmodelling), and the contiguity of the assembly on which the consensus sequences were predicted (Sproul et al. 2023). It is relatively common to use a preexisting repeat library, which may include manually curated TE consensus sequences from one or a few species. However, manual curation of TE consensus sequences is extremely labour-intensive and rarely available for comparative analyses. Our results suggests that uniform repeatmasking provides an efficient compromise for comparative studies, allowing more accurate gene annotations and thus more reliable downstream analyses.

These investigations thus underscore the necessity of annotation frameworks that reduce heterogeneity across species. The uniform annotation pipeline adopted here was overall more successful in standardizing protein length distributions across species compared to the native annotations, while maintaining comparable BUSCO completeness The position overlap of coding sequences shared between uniform and native annotations is ∼80%, which is comparable to a similar comparison of genes detected in different *Bombyx mori* annotations (Han et al. 2024). This type of annotation comparisons has rarely been done, so the generality of this amount of overlap is difficult to assess. While labile gene predictions generally present a caveat, the issue is less grave for gene family evolution analysis. CAFE5 only models gene families present at the phylogenetic root, lineage-specific or *de novo* genes are excluded, minimizing false positives and negatives in orthogroup inference. We conclude that standardization of genome annotations for comparative analyses can be successfully achieved by a common computational pipeline based on a common set of protein homology reference data, on consistently repeatmasked genomes. However, while RNA-seq data can introduce heterogeneity, gene predictions in absence of it can also overestimate gene numbers, which is in turn exacerbated in larger and more repeat-rich genomes that tend to have more fragmented assemblies. It is important to remain aware that any genome annotation represents a *prediction* that can be heavily influenced by methods and reference data rather than a definitive catalogue of genes. All annotation methods generate some false positives and negatives, and the choice of pipeline can therefore influence downstream comparative analyses.

### Genome size and repetitive elements as correlates of gene family evolution

Our results reveal how even closely related species can show large variation in the number of predicted genes, which is mostly due to changes in gene family sizes rather than novel genes (Fig 3B). To understand such variation in its wider genomic context, we tested the impact of GS and repeat abundance on gene family proliferation. The overall repeat abundance is highly variable and different repeat classes dominate the genomes of different species. While highly repeat-rich genomes tend to have a more similar distribution of repeats, in repeat-poor species the repeats are localized close to centromeric regions. Moreover, the overall repeat abundance correlates positively with genome size, in line with other insect taxa, suggesting that repeat accumulation is a major driver of genome expansion. There is no clear phylogenetic signature of GS or RE variation, suggesting that the genome size and its repeat content evolve rapidly and in a lineage-specific way.

TEs can facilitate duplications both directly through mechanisms such as retrotransposition or non-allelic homologous recombination between repetitive sequences, and indirectly, by promoting local structural rearrangements that increase copy number variation. Repeat-rich genomes may therefore experience higher rates of gene copy gain. In addition, TE activity and other repeat-mediated structural variation can also affect regulatory regions to enable expression and functional divergence of paralogs, which can allow duplicates to persist. Together, these processes could create a permissive genomic environment where both structural and regulatory innovation are more frequent.

Here, we detected a positive association between genome-wide repeat abundance and gene family sizes, in line with the fact that repetitive elements contribute to the genomic substrate for gene duplication. Our comparative approach reveals that on average more repeat-rich genomes are prone to harbor greater gene family expansions. However, there was no genome-wide association between any particular TE class and gene family size. Instead, we discovered individual orthogroups that are significantly correlated with more simple sequence repeats such as satellites and low complexity regions (Fig. S13 B and D), the latter of which is correlated with an odorant binding protein. Lack of stronger genome-wide patterns can be expected since repeats may also accumulate in gene-poor regions, where they may be less important for gene family expansions, thus distorting genome-wide signals across the phylogeny (see Figures S23, S25, S29 and S30). On a more local level we do however detect that TE landscape matters. We find evidence that genes in expanding gene families are significantly associated with elevated mean abundance of LINEs and LTRs close to the genes in all species, and DNA transposons in most, although the support for the latter is much weaker (Figure 4 and Figures. S46 to S103). It is important to consider that LINEs and LTRs are Class I TEs that do not maintain introns while DNA transposons are Class II TEs that keep introns. Our finding that significantly rapidly evolving gene families have a higher proportion of intron-less genes (Fig. S15), support that retrotransposition is an important agent of gene family proliferation in beetles. This is not a general rule, since DNA transposons were elevated in *B. siliquastri, T. molitor, R. ferrugineus* and *Z. morio,* with Helitrons (also Class II) also being enriched in the latter two. Our findings thus highlight that repetitive elements are associated with gene family size most prominently on a local scale, and TEs of both classes can facilitate gene duplications, with Class I being more important overall. We conclude that repetitive elements are putative facilitators of gene family expansion and long-term genomic innovation in beetles.

We detected a positive correlation between genome size and average gene family size (Fig. 3D), indicating that larger genomes tend to harbor more extensive repertoires of homologous genes. Because much of the variation in genome size across beetles stems from repetitive DNA (Fig.3A), this relationship likely arises through indirect effects of repeat-mediated duplications and structural rearrangements that accompany genome expansion. Larger genomes may also be more permissive to the retention of duplicates if relaxed purifying selection, reduced recombination, or lower gene density mitigate the costs of redundancy. In such genomes, additional copies may persist long enough to diverge functionally, contributing to lineage-specific innovation. Larger genomes can also provide a greater substrate for transcriptional networks that facilitate retaining duplicates by sub- and neofunctionalization. It is interesting to note how some studies have discovered a positive relationship between GS variation and fitness-related traits both between (Arnqvist et al. 2015) and within species (Boman and Arnqvist 2023)(Stelzer et al. 2021). This seeming contradiction of larger genomes being associated with higher TE content *and* improved fitness could be understood by considering gene duplications. There are now numerous examples how gene duplications can improve fitness by generating functional diversity ((Dort et al. 2024) (Karunarathne et al. 2025) (Tralamazza et al. 2024) (Wei et al. 2025)). Taken together, our findings suggest that genome size variation is not only affected by commonly considered neutral processes, but also correlates with adaptive capacity through gene family proliferations.

### Functional implications: rapid expansions in chemosensory and detoxification genes

Nearly 500 rapidly evolving gene families detected here provide insight into the functional diversification of Polyphaga beetles. These families are enriched in chemosensory receptors and odorant binding proteins, pheromone synthesis proteins, detoxification enzymes, and other genes linked to sexual reproduction and immunity. For example, the most prominent functional cluster of detoxification genes includes sub-families of Cytochrome P450 (CYP), with significant expansions in *Elateroidea* (*I. luminosus* and *P. pyralis*) and *Tenebrionoidea* (*T. castaneum, Z. morio* and *T. molitor*) (Fig. 5 C and D). Aldehyde oxidase detoxification enzymes have expanded in these groups as well, and contribute to pesticide resistance (Zhu et al. 2013). Larvae of the “superworm” *Zophobas morio* can digest plastic via polyethylene oxidization by cytochrome P450 and the lipid metabolism pathway (both insect-encoded and not part of the gut microbiome) (Kim et al. 2024). Furthermore, fireflies (*Elateriodea, I. luminosus* and *P. pyralis*) achieve fluorescence through luciferase which has evolved from a gene duplication of acyl-CoA synthetase (Fallon et al. 2018b), and we identify significantly rapidly evolving orthogroups that show large expansions in the relevant species (Fig. S19). Finally, several orthogroups related to host and mate identification show rapid evolution (Fig S20 and S21). All of these gene families fall into the broad categories of host adaptation and communication and are repeatedly implicated in ecological adaptation and radiation in insects (Singh et al. 2025) (Fang et al. 2009). Expansion in these families likely underlies the capacity of beetles to exploit diverse ecological niches, including novel host plants and chemically defended environments. Similar expansions in chemosensory and detoxification families have been observed in Lepidoptera (Dort et al. 2024)(Zhao et al. 2017) and other arthropods (Sánchez-Gracia et al. 2009), suggesting convergent evolutionary strategies in the adaptation of phytophagous insects. The combination of structural flexibility provided by a repeat-rich genome and ecological selection for chemical diversification may thus be a key factor in the extraordinary success of Coleoptera.

## Methods

### Comparison of genome annotation strategies

First, we compared heterogenous ‘native’ annotations of each species to uniform annotations, created using the BRAKER3 pipeline, which utilizes a combination of *ab initio* and homology-based approaches. The uniform annotation strategy included the following steps.1) Repeats in each assembly were modelled with RepeatModeller v.2.0.4 (Flynn et al. 2019) and masked with RepeatMasker v.4.1.5 (Smit et al. 2015), to annotate the repeats in a uniform way and to create species-specific libraries of repeat consensus sequences for each species for downstream analyses. 2) The repeatmasked assemblies were annotated using the BRAKER3 singularity container (Hoff et al. 2019) (Brůna et al. 2020) (Buchfink et al. 2014) (Gotoh 2008) (Iwata and Gotoh 2012) (Stanke et al. 2008) (Stanke et al. 2006) (Gabriel et al. 2021) and using protein reference data from the orthoDB Arthropoda v11 (Kuznetsov et al. 2023). 3) Each annotation was then filtered to resolve overlapping gene models and to retain only the longest isoform using AGAT v.1.3.2 (Dainat J. 2022). The protein sequences were extracted from the processed annotations using gffread v.0.12.7 (Pertea and Pertea 2020) and emboss v.6.6.0 (Rice et al. 2000). Protein sequences with an incorrect start or stop codon were removed. 4) The transcripts were further filtered for TEs that escaped masking, by removing proteins that have a BLASTp v.2.13.0 (Camacho et al. 2009) hit with species-specific repeat consensus sequences identified by RepeatModeller (e-value < e-10 and sequence identity > 90%), following Hassan et al. (Hassan and Adelson 2023). Additionally, we repeated this hybrid annotation pipeline on all assemblies but without the steps of unified repeat handling, to demonstrate the importance of consistent repeat-modelling and -masking for gene prediction. The overlap of coding regions between different annotations (native and uniform) on the same assembly was computed using BEDtools v.2.31.1 *intersect* (Quinlan and Hall 2010). To compare each annotation strategy, we used BUSCO v.5.5.0 (Manni et al. 2021) (insecta_odb10) database for evaluating completeness of annotation of conserved proteins. We also investigated overlap of detected genes between the annotations, based on the position information of the gene models. Both native and uniform annotations for each species are based on the exact same assembly, therefore we can compare which sequence positions are annotated in each method by position using BEDtools. We compare the overlap in both directions (native annotation query to uniform annotation reference, and vice versa), and consider three possible overlap categories: no overlap, 1-to-1, or 1-to-many (see Fig. 2D for a schematic representation). Furthermore, we use custom python scripts to compare the proportion of single-exon genes (intronless genes) and protein length distributions.

Second, we tested the impact of population origin of RNAseq data on gene detection when using evidence-based hybrid genome annotation approach, by annotating the same genome assembly, of *Callosobruchus maculatus* bruchid seed beetle, with three alternative sources of RNA-seq data that vary by population origin but were otherwise comparable (the same adult life history stage and overlapping sets of tissues). The genome assembly (Kaufmann et al. 2024) is from a population collected from Lomé, Togo (west Africa), and the RNA-seq datasets were obtained either from the same population (Lomé (Kaufmann et al. 2024)), or from two geographically distinct populations (Nigeria (Lu et al. 2024) and South India (Sayadi et al. 2016), see SI for accession numbers). The annotation pipeline is described below and in Fig. 1A (also see extended flowchart in fig. SR3) and was run once for each RNA-seq dataset, resulting in three uniform protein sets.

### Analyses of genome size, repeat content and gene family expansions

The protein sets generated by the unified annotation strategy were used for all downstream analyses. First, orthogroups were identified with OrthoFinder v. 2.3.3 (Emms and Kelly 2019), which also provides a phylogenetic tree through Fasttree, and orthogroups are filtered down to only ancestral orthogroups that are inferred to be present at the root of the tree due to their presence in the outgroup *D. melanogaster.* CAFE5 v.5.0 (Mendes et al. 2021) was used to identify significantly rapidly evolving orthogroups across the phylogeny. The significantly rapidly evolving orthogroups were functionally annotated using information from *Drosophila melanogaster* orthogroup members by automatically extracting functional information from the Flybase API. To detect functionally related groups of rapidly evolving gene families, we clustered the *D. melanogaster* orthologs from each significant gene family into functional categories using DAVID Bioinformatics (Sherman et al. 2022)([CSL STYLE ERROR: reference with no printed form.]), resulting in 36 clusters of rapidly evolving gene families. We have then further manually curated this clustering based on the above-mentioned functional annotation to focus on functional categories that are known to be important in other insect orders, such as processes involved in pheromone communication and host compatibility. See the full functional annotations in Supplementary File 1. We run CAFE5 20 times to ensure convergence and only consider orthogroups as significant when they were significant in every CAFE5 run.

In order to estimate associations between repeat content or genome size with the size of significantly rapidly evolving gene families within each species, we obtained total repeat abundances from RepeatMasker output. Genome size (GS) was estimated from the assembly size, which has been shown correlate closely with flow cytometry estimates (Elliott and Gregory 2015) (Pflug et al. 2020). To account for phylogenetic relationships, we computed phylogenetically independent contrasts for all variables in the statistical analysis using a custom python implementation following the method by Felsenstein (Felsenstein 1985). We calculate correlation coefficients via Spearman’s ranked order correlation (Spearman 1904) as implemented in SciPy (Virtanen et al. 2020) and use Benjamini-Hochberg correction for multiple testing to assess the significant correlation of individual orthogroups. Further, we calculate the mean Spearman’s ρ and 95% standard error of the mean for all orthogroups in each comparison to assess if it is significantly different from 0.

We assess the repeat landscape around transcripts that are part of significantly rapidly *expanding* gene families in each species by only considering transcripts when their associated gene family has at least two members in the species of interest. We use the software package ReVis v.2.3.2 (Trabert 2026) and the repeat annotation generated by RepeatMasker to test for enrichment of certain repeat categories around transcripts that are part of expanding gene families.

## Data availability

Annotations and assemblies of all selected species were downloaded from the NCBI. All code for the analyses can be found here: https://github.com/milena-t/PhD_chapter1.

## Author contributions

MT and EI conceived and designed the study. MT performed the analyses and generated figures. MT and EI wrote the manuscript. JB contributed to conceptual and methodological development and provided critical revisions. EI supervised the project and secured funding. All authors approved the final version of the manuscript

## Acknowledgements

We would like to acknowledge Martin Pippel at the National Bioinformatics Infrastructure Sweden at SciLifeLab for bioinformatics advice. The computations were enabled by resources in project UPPMAX 2026/1-8 provided by the National Academic Infrastructure for Supercomputing in Sweden (NAISS) at UPPMAX. This work was supported by a grant from the Swedish Research Council (Vetenskapsrådet) (grant no. 2023-04869) to EI, JB acknowledges support from The Birgitta Sintring Foundation, Lennanders Foundation and the Swedish Research Council (Vetenskapsrådet) (grant no. 2025-00450).

